# An automated confocal micro-extensometer enables in vivo quantification of mechanical properties with cellular resolution

**DOI:** 10.1101/183533

**Authors:** Sarah Robinson, Michal Huflejt, Pierre Barbier de Reuille, Siobhan A. Braybrook, Martine Schorderet, Didier Reinhardt, Cris Kuhlemeier

## Abstract

How complex developmental-genetic networks are translated into organs with specific 3D shapes remains an open question. This question is particularly challenging because the elaboration of specific shapes is in essence a question of mechanics. In plants, this means how the genetic circuitry affects the cell wall. The mechanical properties of the wall and their spatial variation are the key factors controlling morphogenesis in plants. However, these properties are difficult to measure and investigating their relation to genetic regulation is particularly challenging. To measure spatial variation of mechanical properties, one must determine the deformation of a tissue in response to a known force with cellular resolution. Here we present an automated confocal micro-extensometer (ACME), which greatly expands the scope of existing methods for measuring mechanical properties. Unlike classical extensometers, ACME is mounted on a confocal microscope and utilizes confocal images to compute the deformation of the tissue directly from biological markers, thus providing cellular scale information and improved accuracy. ACME is suitable for measuring the mechanical responses in live tissue. As a proof of concept we demonstrate that the plant hormone gibberellic acid induces a spatial gradient in mechanical properties along the length of the Arabidopsis hypocotyl.

**Terms:**

Stress: is the force acting on the material per unit area.
Strain: the relative increase in length of the material, and can be expressed as a percentage change in length.
Mechanical properties: describe the stress-strain relationship for a material. If we apply the same force to a material that is twice as thick/stiff? it will deform half as much, if the material is otherwise the same.
Elastic: elastic materials deform instantly and reversibly.
Creep: a time-dependent irreversible strain that occurs when a constant force is applied and maintained. Creep is measured using creep tests. A force is applied and maintained for a period of time. The force is removed to reveal the reversible and irreversible deformation.

## Introduction

Understanding how gene activities are translated into shapes is still a major challenge. The key to deciphering this process is to have a better insight into the role of mechanics (Moulia et al., 2011). Growth occurs by the yielding of the cell wall to stress (Lockhart, 1965) and the direction of expansion is controlled by the relative properties of the tissue in the different directions. Altering these properties will lead to the formation of different shapes (Coen et al., 2004; Kuchen et al., 2012; Green et al., 2010). At the cell wall scale these properties are primarily determined by the orientation of cellulose fibers (Green, 1962; Probine and Preston, 1961), which are deposited by cellulose synthase complexes that track along the microtubules (Paredez et al., 2006). Localized changes in the expression of cell wall modifying proteins will alter the wall’s ability to expand and result in differential tissue deformation and, therefore, control morphogenesis (Fleming et al., 1997; Pien et al., 2001; Peaucelle et al., 2008).

Morphogenesis is also influenced by mechanical feedback. Plants are known to sense mechanical stress such as wind, gravity, bending, even touch, and alter their growth accordingly (Bastien et al., 2013; Band et al., 2012; Chehab et al., 2009; Braam and Braam, 2004; Ditengou et al., 2008; Richter et al., 2009). There is mounting evidence that plants can also sense internal mechanical stress with microtubule orientation being reported to correlate with stress patterns. Such internal tissue stresses typically arise due to the geometry of the tissue (Dumais and Steele, 2000; Hamant et al., 2008), different properties of cell layers (Peters and Tomos, 1996; Hejnowicz, Z, and Sievers, 1995; Kutschera and Niklas, 2007), or differential growth across a tissue (Coen and Rebocho, 2016); all of which are in turn controlled by mechanical properties. These stresses have also been proposed to influence morphology directly by causing the tissue to buckle (Eldridge et al., 2016; Green et al., 2010, Green et al., 1999).

In order to fully explore how local wall properties translate into specific shapes and how they interact with gene regulatory networks, there is a need for techniques that enable mechanical properties to be quantified in developing tissues and responses to mechanical stress to be observed with high spatial resolution. Mechanical properties are intrinsically difficult to measure as force can only be measured by its impact on an object. The mechanical properties of a material describe how it deforms when a force is applied; formally they describe the relationship between stress (force/cross sectional area) and strain (relative change in length). Mechanical tests, therefore, depend upon precisely applying either a force or a deformation to a tissue and measuring the other property.

Many methods are available for measuring tissue level mechanics in large samples: including extensometers. The classical extensometer setup involves clamping the sample, then applying a calibrated weight and measuring the deformation. The samples are typically centimeters in length and deformations are measured in the order of millimeters per hour. Extensometers were used extensively and most notably in the discovery of the cell wall modifying protein expansin (McQueen-Mason et al., 1992). Expansins were found to increase cell wall creep: a time-dependent irreversible strain that occurs when a constant force is applied. The irreversible component is calculated by removing the force and measuring the deformation that remains. This type of test is thought to best reflect the action of turgor on the cell wall. These experiments were conducted on dead tissue so that water movement would not be an issue, and were boiled to inactivate endogenous enzymes and proteins.

Driven by the need to study mechanical properties in the smaller developing tissues of Arabidopsis nano- and micro-indentation techniques have been adapted for this purpose. All of these methods involve indenting the tissue and measuring the force required to do so. Atomic force microscopy (AFM) is such a technique and was used to identify spatial differences in the cell wall properties in the shoot apical meristem (Milani et al., 2011). These experiments were performed on plasmolysed tissue and involved very rapid indentations (30-80μm s-1) of 40-100nm in depth using a tip with a radius of 10-40nm. These type of measurements provided very high resolution local information. Indenting with larger probes (2-11μm) has been proposed to provide information on inner layers (Peaucelle et al., 2011; Braybrook and Peaucelle, 2013), or about the turgor pressure of the cells (Routier-Kierzkowska et al., 2012; Weber et al., 2015). Extracting cell wall or turgor pressure measurements from indentations requires sophisticated models that take into account the relative contribution of the geometry, cell wall thickness, etc. Measurements are also made perpendicular to the main direction of growth. This is appropriate for the study of the pectin matrix or turgor as they are isotropic, however, the other structural cell wall components such as cellulose fibers are highly anisotropic and it is less clear how this information should be interpreted (Cosgrove, 2015).

AFM and extensometers provide very different information and operate at very different scales. Here we propose a new technology; an automated confocal micro-extensometer (ACME). ACME can be used to measure mechanical properties and also to apply mechanical stress. Designed to bridge the gap between classical extensometers and AFM, ACME provides tissue and cellular resolution information on the mechanical properties of small growing Arabidopsis tissues. By facilitating mechanical measurements on developing Arabidopsis tissues, we expand the possibility of utilizing the vast array of knowledge and genetic tools that have been developed by the community.

Conceptually, ACME operates like a classical extensometer but is much smaller, fully automated and crucially relies on confocal images for strain computations. Strain is, therefore, computed from features on the tissue itself. This improves the resolution, accuracy and scalability of the measurement. The simultaneous acquisition of images also enables responses to mechanical stress and tissue health to be assessed continuously and in real time. Extensometers have previously been used to study elasticity and creep in Arabidopsis but have been restricted to larger samples (Park and Cosgrove, 2012; Miedes et al., 2013) and were not coupled to a confocal microscope. Extensometers are naturally easier to couple to a confocal microscope as they measure perpendicular to the imaging axis. Indentation methods by contrast obscure the image, and must be used sequentially, (Milani et al., 2014; Louveaux et al., 2016).

We demonstrate the usefulness of ACME by investigating the response of light grown Arabidopsis seedlings to the growth hormone gibberellic acid (GA). GA is known to promote a burst of growth in light-grown Arabidopsis hypocotyls (Sauret-Güeto et al., 2012). Studies in other species have demonstrated that GA acts to regulate growth via changes in the cell wall (Adams et al., 1975; Stuart and Jones, 1977; Cosgrove and Sovonick-Dunford, 1989). Here we study the mechanical changes that occur during the response to GA in the tiny light-growth hypocotyl of Arabidopsis thaliana with cellular resolution.

## Introduction to ACME

In order to be able to measure a range of material properties, at different time-scales, and with cellular resolution: we developed a miniature extensometer which is mounted on a confocal microscope (Figure 1A-B). It is tailored for use on small samples, such as Arabidopsis seedlings but can be adapted for larger samples and used in combination with other imaging systems. ACME is easily assembled from a combination of commercially available parts, and custom parts which can be 3D printed (designs included) or cut from sheet aluminum (see methods).

**Figure 1.**
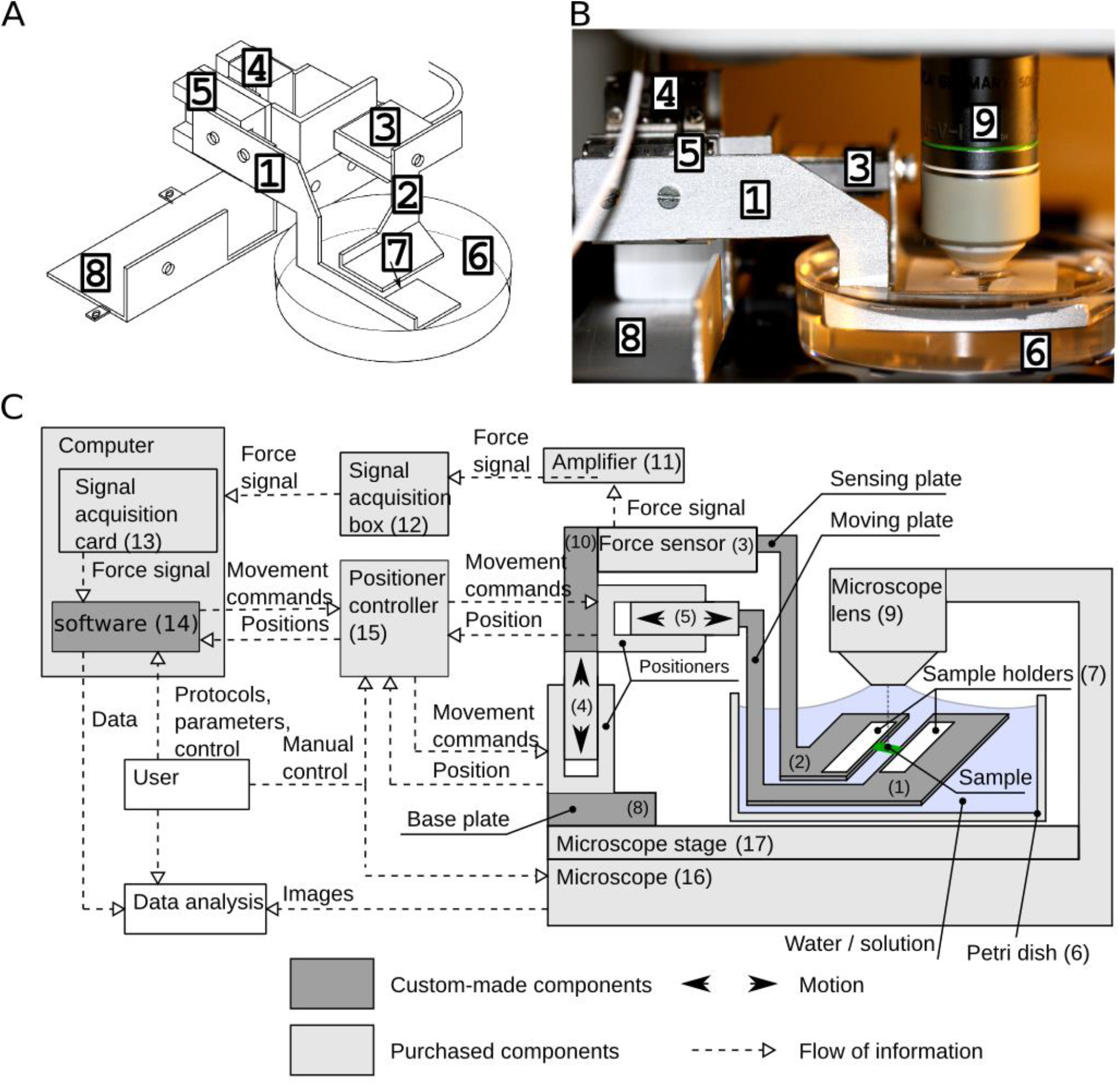
The Automated Confocal Micro-Extensometer (ACME) setup. **(A)** A diagram of ACME. **(B)** Photograph of ACME mounted on Leica SP5. **(C)** Diagram of the control of ACME. The panels show: **(1)** the moving arm; **(2)** the measuring arm; **(3)** the force sensor (Futek LSB200 10g load cell); **(4,5)** the nano-positioner (SLC1720; SmarAct GmbH); **(6)** the dish for solution; **(7)** the sample mounting position; **(8)** the confocal mount; **(9)** a 20x confocal dipping objective. **(11)** amplifier (Futek CSG110) Futek Inc.); **(12)** SCB68 signal acquisition box; **(13)** computer-based signal acquisition system (NI6221PCI); **(14)** custom-made software and SmarAct controller software library; **(15)** SmarAct MCS3D (SmarAct GmbH) controller); ACME is a combination of custom and purchases components shown in dark and light gray. **(1, 2, 8, 10)** Custom made aluminium parts; **(9, 13, 16, 17)** Microscope.

Like a classical extensometer it enables measurements to be made parallel to the direction of growth. The sample is manipulated using a robotic nanopositioner (Figure 1A-C, label 5), which enables very small movements to be made with high accuracy (better than 50nm). ACME measures forces using a load cell, so that the force can be measured instantly and directly, rather than having to be computed. For this study we used a 10g load cell to enable forces to be measured within the physiological range for a light grown Arabidopsis hypocotyl (Figure 1A-C, label 3) and with low noise (<100μN per hour (Figure S1)). We typically measured tensile forces between 1mN and 10mN (0.1-1gram). Custom software connects the positioner and the force measurement in a feedback loop (Figure 1C label 14) so that a stable force can be maintained. In order to provide a positive user experience, full flexibility in the type of experiment and to ensure reproducibility, the user can control ACME via small protocols that are defined in the parameter file. The user can specify a force that should be applied or a deformation, and the duration for which it should be maintained. Any number or combination of forces or deformations can be specified. ACME will then perform these steps and record the force and position continuously (Figure 2A-F). The user can also determine the tolerance around the target force, and the step size of the robot. The ability to perform a wide range of experiment types is important as biological materials are heterogeneous and their properties cannot be defined by a single experiment type.

**Figure 2.**
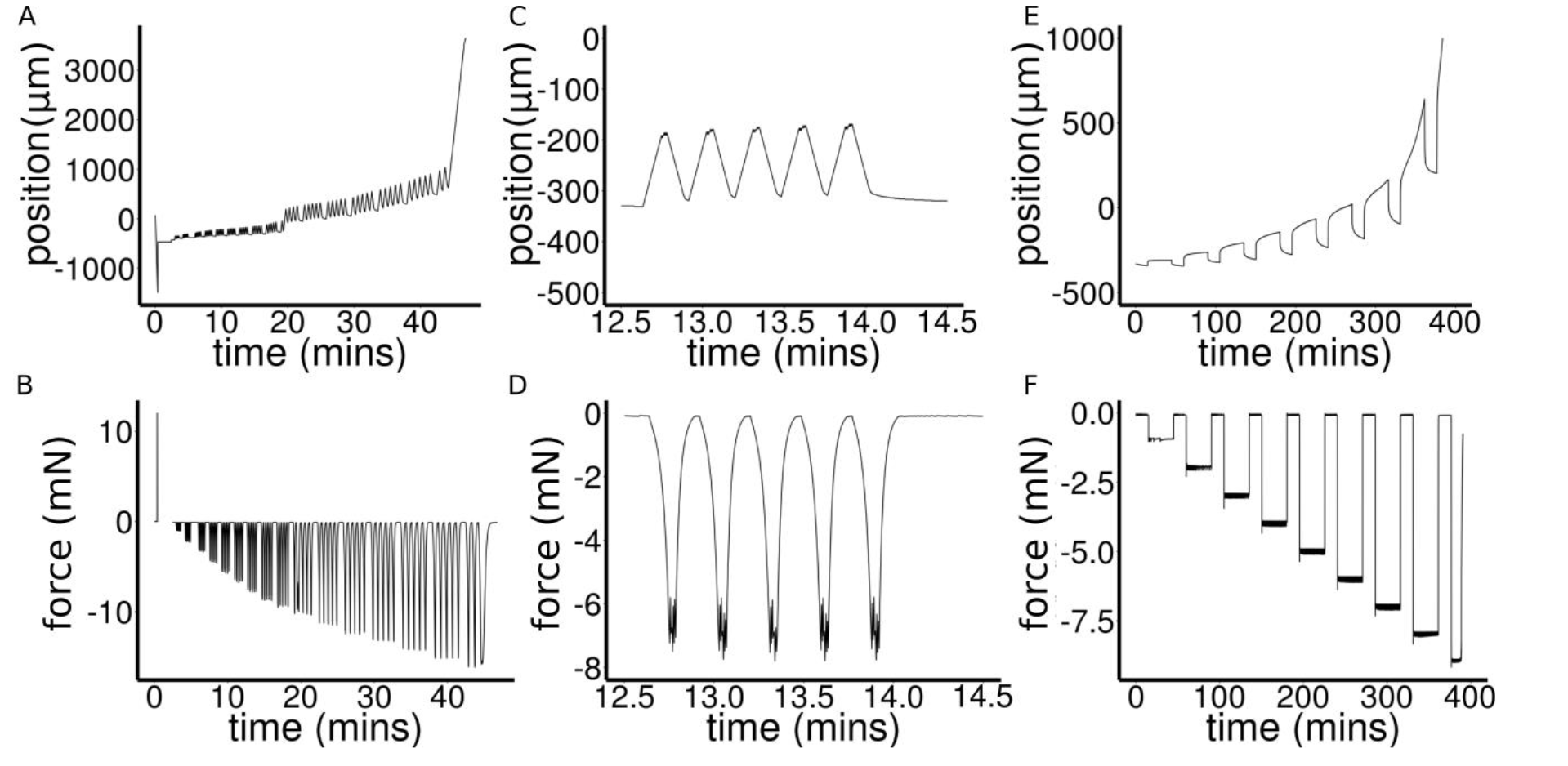
Examples of ACME experiments. **(A, C, E)** robot position against time and **(B, D, F)** force against time. **(A-B)** Oscillation experiments. Initially there is a calibration step then the experiment begins. The force is applied in groups of 5 oscillations with a 30 second pause at 0 force in between. At the end the sample detaches and the experiment stops. Some slippage is apparent at t= 20 min, which is why material coordinates are used for the analysis of strain. **(CD)** A subset of experiment in A-B. **(E-F)** Multi-creep curves showing sample being repeatedly loaded and held at a fixed force before being returned to 0 force.

ACME is sufficiently small and light to enable mounting onto a confocal stage without obscuring the image or impacting the microscope function (Figure 1B label 9). A custom base plate for secure attachment to the z-stage (Figure 1A-C label 8), enables simultaneous image acquisition and mechanical measurements to be made. The images are used to compute strain/deformation more accurately compared to using the robot co-ordinates, which are sensitive to any sample movement. The images also enable responses to stress or strain to be observed at a cellular level.

### Measuring mechanical changes that occur in light grown hypocotyls

We demonstrate the usefulness of ACME by using it to measure the mechanical changes that accompany early growth in light grown hypocotyls. We characterised the growth of seedlings in the presence of either GA or the GA-biosynthesis inhibitor uniconazole compared to control conditions. GA treated seedlings were much longer five days after stratification (DAS) compared to the control or uniconazole treated seedlings (Figure 3A). The majority of this growth occurred between 2DAS and 3DAS. Uniconazole prevents germination, so seedlings were transferred to the uniconazole at 2DAS which suppressed subsequent elongation. The inhibitor was used rather than GA mutants as the mutants often had a disrupted geometry. The difference in hypocotyl length is significant (<0.00001) between the GA and control treatments at 3-5DAS but not at 2DAS. The inhibitor treated samples are significantly smaller by 5DAS.

**Figure 3.**
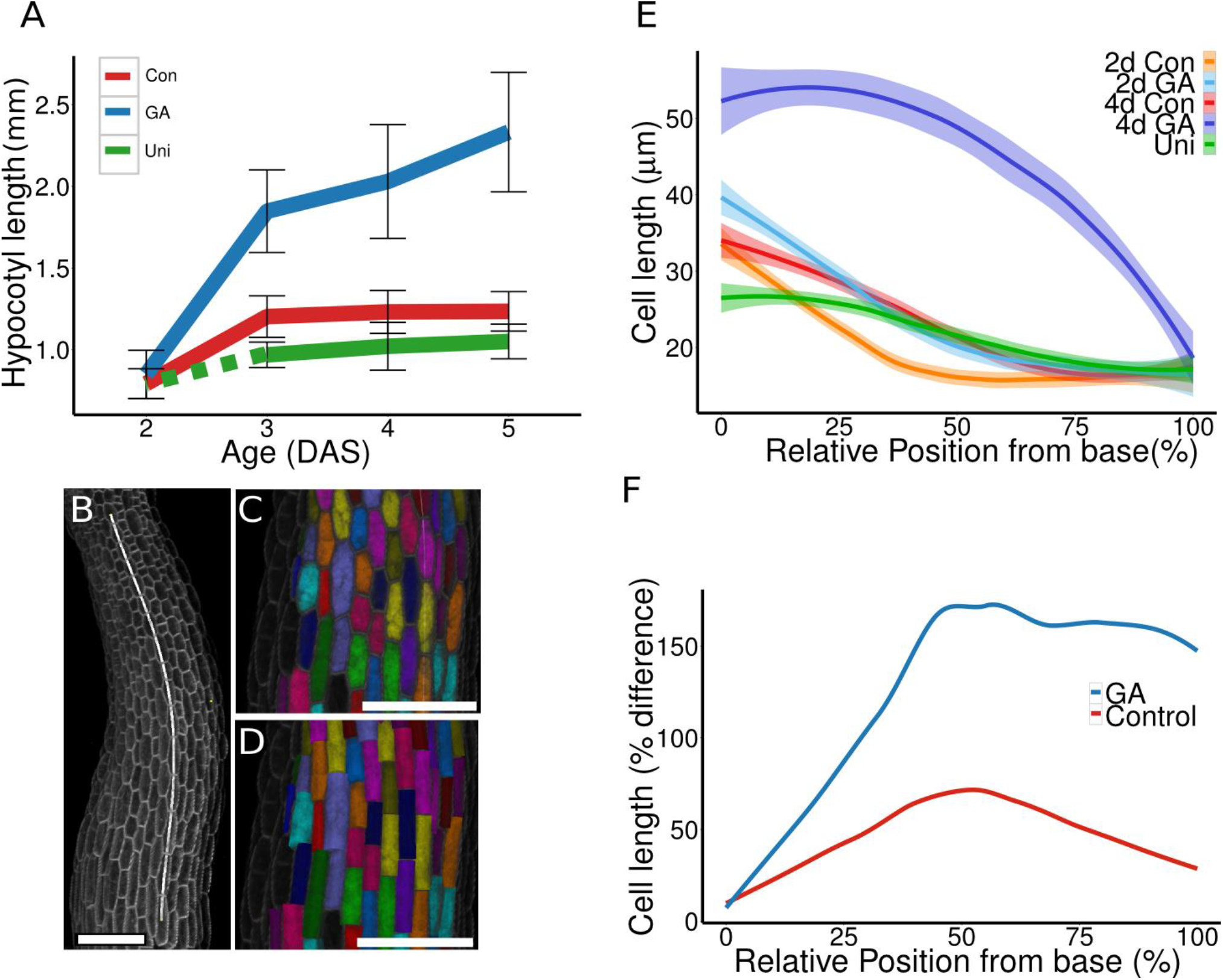
The growth response to GA is under tight spatial control. **(A)** Lengths of hypocotyls grown in control conditions (Con) in the presence of GA or after being moved to uniconazole (Uni) (dashed line indicates seedlings approximate length before being moved). Error bars show standard deviation. **(B)** A file of cells was selected and a Bézier curve of order 5 was fitted. This enables distances to be displayed in terms of distance along the hypocotyl. **(C)** The cells are segmented in 3D and assigned a random label and colour to identify them. Cells that were not segmented well were deleted or manually corrected. The volume of the cells was extracted directly from the segmented image. **(D)** Shows the same cells as in (C) but showing the cells approximated as cylinders. In order to extract the cell length and mean radius we must extract the principal components of the shape. The cell length is taken to be the PC1, the average of PC2 and 3 is taken for the width. The three PCs can be visualized as a cylinder, enabling badly represented cells to be manually removed. The high similarity between C and D shows that the approximation is good. (**B-D**) Scale bar is 100 μm. **(E)** Using the PC1 data cell length was computed as a function of the distance from the base of the hypocotyl. The 99% confidence interval of the mean is shown. **(F)** The relative change in cell length that occurs between 2 and 4DAS for control and GA treated seedlings is shown against relative distance from the base of the hypocotyl. 0 is the base of the hypocotyl just above the root and 100% is the top of the hypocotyl just below the cotyledons. The increase in length is greater in the GA treated seedlings and is not uniform along the length of the hypocotyl and is much greater in the top half of the hypocotyl. GA data is from 1500 cells from 10 seedlings, and control data is from 1000 cells from 5 seedlings.

Epidermal cell size was measured in 3D using the image analysis software MorphoGraphX (Barbier de Reuille et al., 2015) (Figure 3B, C). Distance along the hypocotyl was determined by fitting a Bézier curve (Figure 3B), and referring to cells position along the curve. The volume of epidermal cells was extracted from the segmented volume (Figure 3C). We compared the volume of cells from the different treatments at different relative distances along the hypocotyl (Figure 3E). At 2DAS, when hypocotyl height was comparable, cell volume was not different between control and GA treated hypocotyls; by 4DAS the volume of cells was larger in the GA treated plants compared to the control. In order to compare the length of the cells, this information must be extracted from the segmented images; this was done using principal component analysis (PCA). PCA enables us to extract the dimensions of the cells resulting in the cells being approximated as cylinders (Figure 3D). PC1 corresponds to the length of the cells. By comparing the length of the cells in hypocotyls at 2 and 4DAS we determined that all of the cells expanded in length (Figure 3E). We computed the change in average cell length from 2DAS to 4DAS by performing local polynomial regression fitting. The GA treated and control seedlings do not expand uniformly along their length (Figure 3F). In the GA treated plants (blue line) the increase in cell length was larger in the middle and upper parts of the hypocotyl while in control seedlings (red line) it occurred in the middle of the hypocotyl.

### High throughput imaging to determine tissue elasticity in the direction of growth

We used ACME to determine if GA treatment altered the elastic properties of the cell wall. Elastic materials deform instantly, and upon unloading instantly return to their original size. By measuring the instantaneous deformation when a force is applied we can compare the elastic properties of tissues. For this type of test the samples were flash frozen, and thawed as in (Durachko and Cosgrove, 2009) so that the water movement would not be a factor. The sample was then rapidly loaded and unloaded multiple times. Such measurements require accurate strain information. The position information from the positioner includes sample slippage which would lead to an over estimation of the strain. Instead we compute strain from the confocal images using landmarks that can be identified on the hypocotyl. The images were acquired continuously generating thousands of images for a single experiment. We developed software to compute the strain from these images. The user must select two regions of interest in the image, usually a cell junction at either end of the sample, or a dot added to the sample. The software then tracks these images and extracts the coordinates (see methods). From these coordinates the oscillations are identified and the strain is computed using R scripts (supplemental file ExtractingOsc.R and oscillations.R).

We compared the strain measured at 5mN of force in samples of different ages and treatments. We found that strain (Figure 4A) was significantly higher in the GA treated hypocotyls (p<0.05) and was lower but not significant in uniconazole treated hypocotyls when compared to control hypocotyls. We also observed that the strain in GA treated hypocotyl was higher in the 2DAS compared to 3DAS samples. These results show a good agreement with the growth data (Figure 3). We conclude that GA increases cell wall elasticity in a temporarily restricted manner. Previous measurements on hypocotyl elasticity were conducted on etiolated samples that are much longer (Park and Cosgrove, 2012). We confirmed that GA increased tissue elasticity by performing AFM experiments (Figure 4B). The apparent stiffness of 2DAS and 3DAS seedlings grown on GA was significantly lower (p<2e-16) than for the control seedlings, consistent with the higher strain measured with ACME.

**Figure 4.**
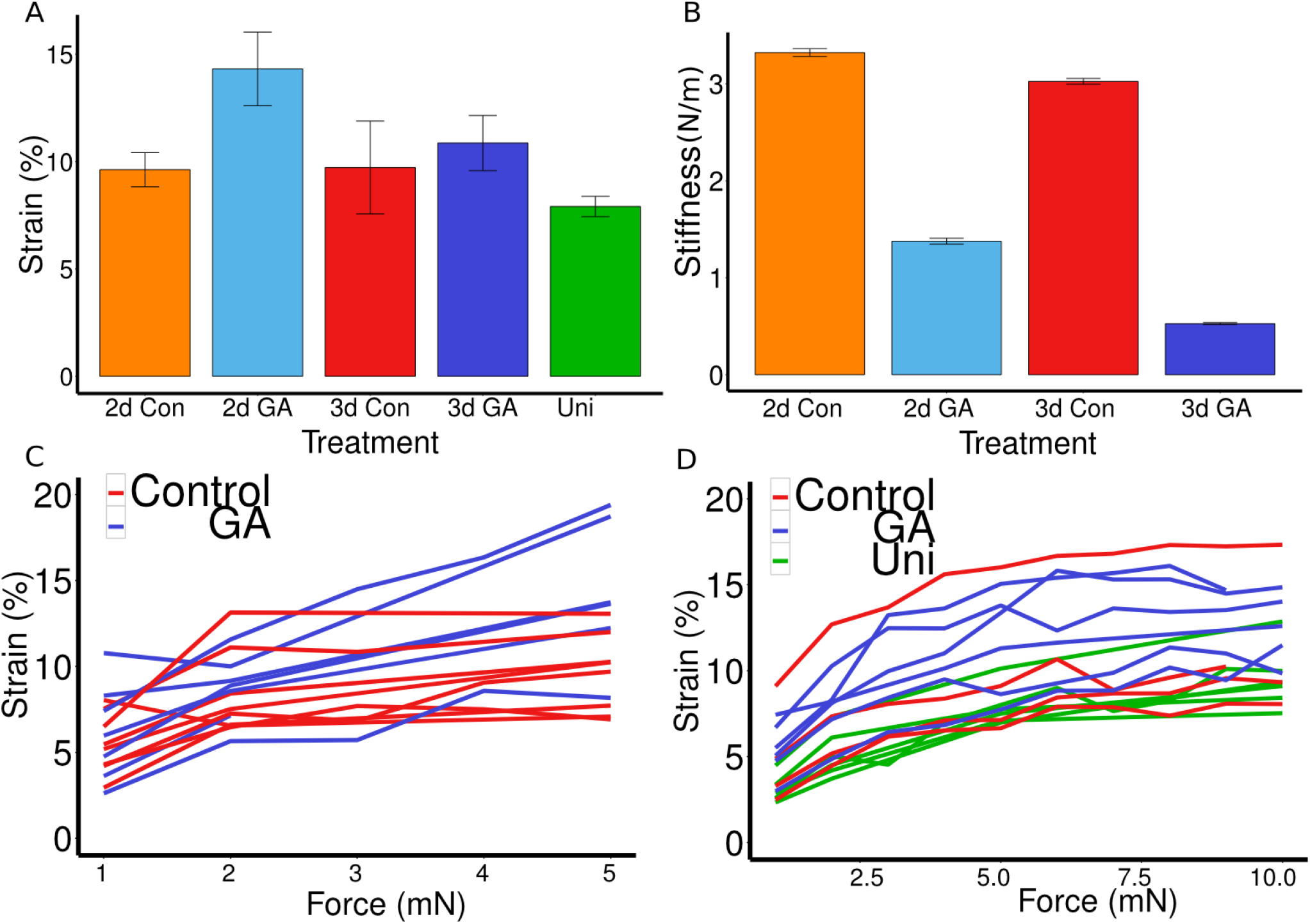
GA increases the elasticity and creep of hypocotyls. **(A)** Hypocotyls were frozen and thawed then subjected to cycles of application and removal of 5mN of force. The average magnitude of strain incurred by 2 and 3DAS, control and GA, and uniconazole treated seedlings is shown, at least 5 seedlings were tested per treatment. Batches of 5-10 oscillations were made. Bright-field images were collected every 645ms. Strain was computed from regions that were tracked in the images using the ACMEtracker software (see methods). The strain for 2d GA differs significantly from the control (p < 0.05, n>6). **(B)** AFM results from 2 and 3 day old seedlings grown in control conditions or in the presence of additional GA. Values show average sample stiffness. For each hypocotyl, three areas of 50x100μm were indented from at least two independent samples. **(C-D)** Average strain was computed from rapid oscillation at a range of forces for 2 and 3DAS seedlings. As above, batches of oscillations were made with 30 seconds at zero force in between forces. Strain was computed from images as described for A.

### Measuring strain stiffening

Materials are usually not linearly elastic (Fung, 1993); meaning that the strain does not correlate in a linear manner with the amount of force applied. The strain may increase more (strain softening) or less (strain stiffening) with increased force. We tested if hypocotyls show strain stiffening behavior by applying a range of forces. The average strain of 5-10 cycles of applying and removing force was plotted against the force applied (Figure 4C,D). The relationship between the two was quantified by fitting linear and Hill-type models to the data and evaluating the fit (Table 1). The linear model had the form Σ=β.F + c and the Hill function had the form Σ =α.F/(λ+F); where Σ is strain and F is force. As both models have two parameters they can be compared by comparing the residual sum of squares (RSS). Although the amount of strain shown by the sample is variable the behaviour of strain against force is more consistent between samples of the same treatment. The stress-strain relationship of 3DAS hypocotyls were all best described by a Hill function which is indicative of a strain-stiffening behaviour. The 2DAS hypocotyls showed reduced strain stiffening when treated with GA. We can conclude that hypocotyls show strain stiffening behavior and this correlates with reduced growth. Strain stiffening in the shoot apical meristem of Arabidopsis has been observed using osmotic treatments (Kierzkowski et al., 2012). Here we were able to demonstrate strain stiffening directly by applying known forces. By using dead tissue for this particular experiment we showed that strain stiffening is a property of the material itself and unlikely to be a consequence of the cell sensing and responding to the mechanical stress. A similar non-linear elastic behaviour has been reported in epidermal peels of maize coleoptiles using an extensometer (Lipchinsky et al 2013). These results suggest that this is a common property of plant tissues.

**Table 1.**
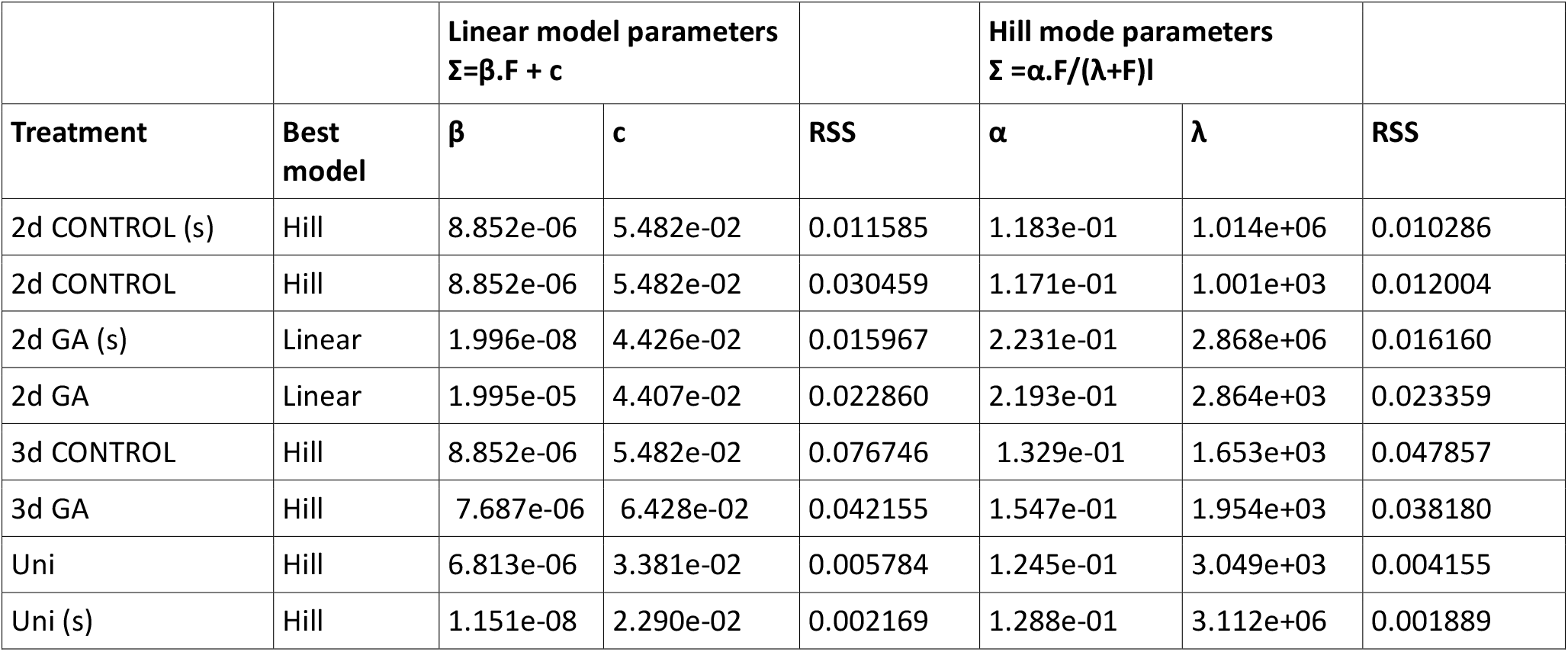
**Comparing the strain stiffening behavior by model fitting.** Model comparison of Linear versus Hill type equation for the oscillation data for the 2 and 3DAS samples for the different treatments. The linear model has the form ***Σ=β.F + c*** and the Hill model has the form ***Σ =α.F/(λ+F);*** Where (Σ=Strain, F = Force). The model fitting is done using the non-linear least square method in R and as both models have two free parameters the model that best fits the data is selected based on it having the smallest residual sum of squares (RSS). Not all data points are present in all datasets for the 2DAS samples so the model was also fit to a subset of the data including only points present in all data sets, and excluding values above 5mN (s). The model chosen is still the same, but the coefficients are different.

### Mechanical measurements on live tissue

In order to be able to utilise the great many fluorescent markers available, we developed a method of attaching the sample to ACME so that it remain alive (see methods). Using this method we were able to maintain hypocotyl samples in the machine for at least 3 hours. The sample health can be evaluated in two ways. If a healthy turgid sample is attached to the device then it will take up water and expand if possible, or generate a pushing force. When the sample is initially mounted the plates are in a fixed position and force will build up. One can then specify to maintain the sample at zero force. The positioner will move the amount that is necessary to maintain zero force. For healthy samples the positioner will keep moving to enable the sample to expand (Figure 5A) and keep the force within the specified tolerance around zero force (Figure 5B). Sample health can also be easily evaluated by observing the cells using the coupling to the confocal microscope (Figure 5C). A live cell is visibly turgid, with the membrane pushed against the cell wall and no visible gaps between the cells. The healthy cells can also express GFP. Counter staining with propidium iodide can also be used to reveal dead cells (Figure 5C arrow). Meanwhile a damaged cell will lose turgor, and stop expressing GFP. When treated with hypo-osmotic mannitol concentrations the cells can be seen to be visible plasmolysed (Figure 5D). As confocal images are collected for the strain calculations, they also serve as an indicator of sample health. Some samples were damaged during the sample mounting and these samples were discarded. Live tissues have different mechanical properties; to compare these we performed short creep tests on turgid, plasmolysed or thawed frozen hypocotyls from 3DAS non-treated seedlings (Figure 5E). Creep tests look at the irreversible deformation of a material when held at a constant force. The sample is typically held at zero force, then a constant force is applied for a period of time, the force is then removed. The permanent deformation is creep, while the recoverable deformation is elastic. This is time dependent and can be observed over long time periods; enabling higher resolution images to be obtained.

**Figure 5.**
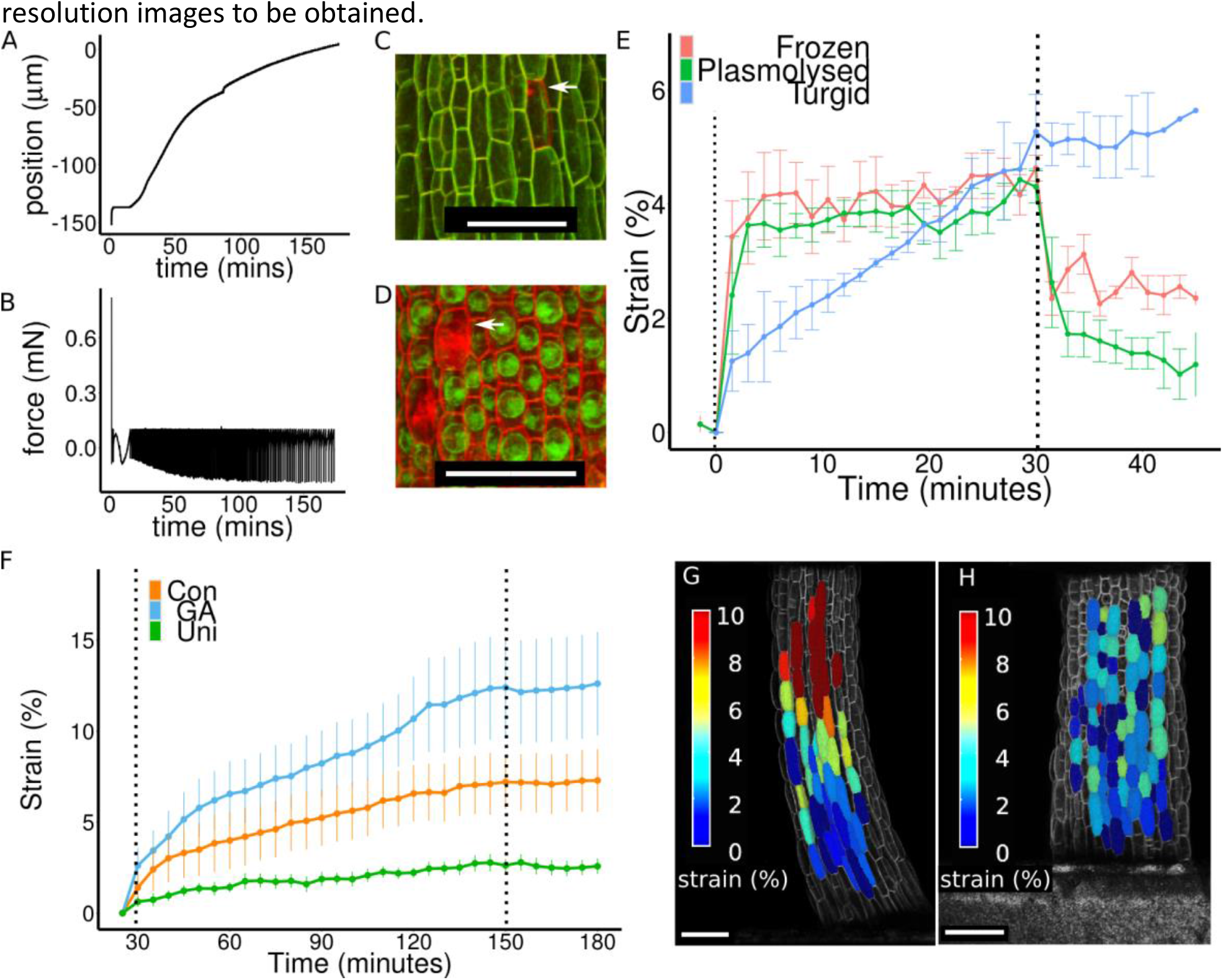
Utilising confocal images to compute mechanical properties. **(A-B)** A live sample will push against ACME as it takes up water and grows. **(A)** The relative position of the plates and **(B)** the force are both recorded. The user can define a holding force of zero, and the threshold of tolerance in this case 0.1mN. When the measured force is outside of defined range, the position changes until the force returns to the within the target range. **(A)** A healthy sample will continue to generate a pushing force and the plate will move to extend the sample and return the force **(B)** to within the range of tolerated forces. **(C-D)** The attachment of ACME to the confocal enables cells to be manually inspected in order to assess their health. **(C)** A healthy sample in water is turgid and the cells express GFP. Staining with propidium iodide (PI) reveals a single dead cell indicated by an arrow. **(D)** By comparison to the turgid cells, when cell are plasmolysed by treating them with 0.5M mannitol in this case or due to poor health this can be easily seen as the protoplasts shrink, dead cells are also visible as they lose GFP expression and they take up PI. **(E)** Creep tests were performed to compare live seedlings, thawed seedlings that had previously been frozen and seedlings plasmolysed with 0.5M mannitol. The samples were loaded with 1mN of force at time 0 and unloaded after 30 minutes. The deformation of the live samples was irreversible, while in the other samples it was partially reversible. At least three seedlings were tested per treatment. Strain was computed from confocal images that were acquired every 1.5minutes. **(F)** Creep tests were performed on live turgid 2DAS seedlings grown in presence of GA, or control conditions and plants grown on uniconazole. GA increased the creep rate and the deformation was irreversible. At least four seedlings were tested per sample. Dashed lines indicate addition and removal of 1mN of force. Strain is shown as a % of deformation. Strain was computed from confocal images that were acquired at a minimum of every 15 minutes. Error bars show standard error. **(G-H)** Creep tests were performed by applying 10mN of force to 3DAS live, turgid samples. Z-stacks were acquired and the images were segmented. Corresponding cells were co-labeled before and after stress was applied. Using PC analysis the length of the cells was computed and used to compute the % strain in length per cell, shown as a heatmap. **(G)** The GA treated sample showed a spatial gradient in % strain along its length. **(H)** The control samples do not show such a gradient in strain. Scale bar = 100μm.

The strain can be measured directly from 2D projections using ImageJ or point tracking software (Kuchen et al., 2012). A force was applied to the sample for 30 minutes then removed. All samples showed a similar amount of strain after 30 minutes; however, in the turgid hypocotyls the strain was more gradual, while in the other samples the deformation was instantaneous (Figure 5E). Upon removal of the force, the turgid sample remained in the deformed configuration and continued to elongate. In both the plasmolysed and thawed frozen samples some of the deformation was recovered, i.e. was reversible. This shows that water movement and/or metabolic processes likely played a role in the non-reversible extension we observed in the live turgid creep tests. Live samples will grow as well as exhibiting creep. We held a live turgid sample at zero force for 3 hours and measured the deformation to be 1% per hour.

### GA increases creep in live tissues

In order to have physiologically relevant data of the response to GA, creep tests were performed on live 2DAS light grown seedlings. The samples were tested just before they would have undergone a burst of GA-induced growth if treated. The sample was maintained at zero force for 30 minutes to allow recovery from the sample preparation. Samples were then subject to 1mN of force for two hours before being returned to zero force for 30 minutes (Figure 5F). Confocal images were acquired every 5-15 minutes and used to track strain. A greater amount of creep was observed in the GA-treated hypocotyls compared to the control, which exhibited a greater amount of creep again compared to the uniconazole-treated hypocotyls (Figure 5F). This confirmed that GA altered the mechanical properties of the cell wall of the seedlings in physiologically relevant conditions. Upon removal of the force there was very little recovery of the initial strain, therefore, the deformation was irreversible within the time frame of this experiment.

### 3D Cellular resolution strain measurements

When force is applied to a sample, the force is equal at any point along its length (Landau and Lifshitz, 1986). Therefore, the difference in strain of the cells, in response to external force can be used to compute their material properties. If a cell strains more than another then it is more elastic. We performed creep tests on live hypocotyls and high resolution images were collected every 5-10 minutes. The images were segmented in 3D and volume changes were computed. PCA was performed to approximate the length, and diameter of cells (average of depth and width). The longitudinal strain of the cells was computed at the end of the creep phase. The cells of the 3DAS GA treated samples showed a much greater amount of strain in the middle and upper parts of the hypocotyl compared to the lower part (Figure 5G). Control seedlings did not show such a gradient (Figure 5H). The gradient in strain along the length of GA treated hypocotyls showed good agreement with the observed growth pattern (Figure 3F). We confirmed that this was not a consequence of there being a difference in the cell wall thickness by analyzing TEM sections taken at different distances along the hypocotyl. If the cell wall was thicker at the base of the hypocotyl this could explain the results without a change in cell wall properties, however, the cell wall was thinnest at the base of the hypocotyl (Figure S2).

## Other tissues

ACME can be applied to tissues other than hypocotyls. Any tissue that is smaller in one dimension compared to the other two, i.e. not spherical, can be measured. We demonstrated this using an Arabidopsis cotyledon (Figure 6). The entire seedling remained intact and the cotyledon was placed between the plates. The sample was healthy before (Figure 6 A,B) and after (Figure 6 C,D) the application of stress as demonstrated by the expression of GFP and the failure of PI to enter the cells. Deformation of the epidermal cells can be visualized by overlaying the before and after image (Figure 6E,F).Further work is needed to interpret the data from cotyledons that have more complex shapes and geometries.

**Figure 6.**
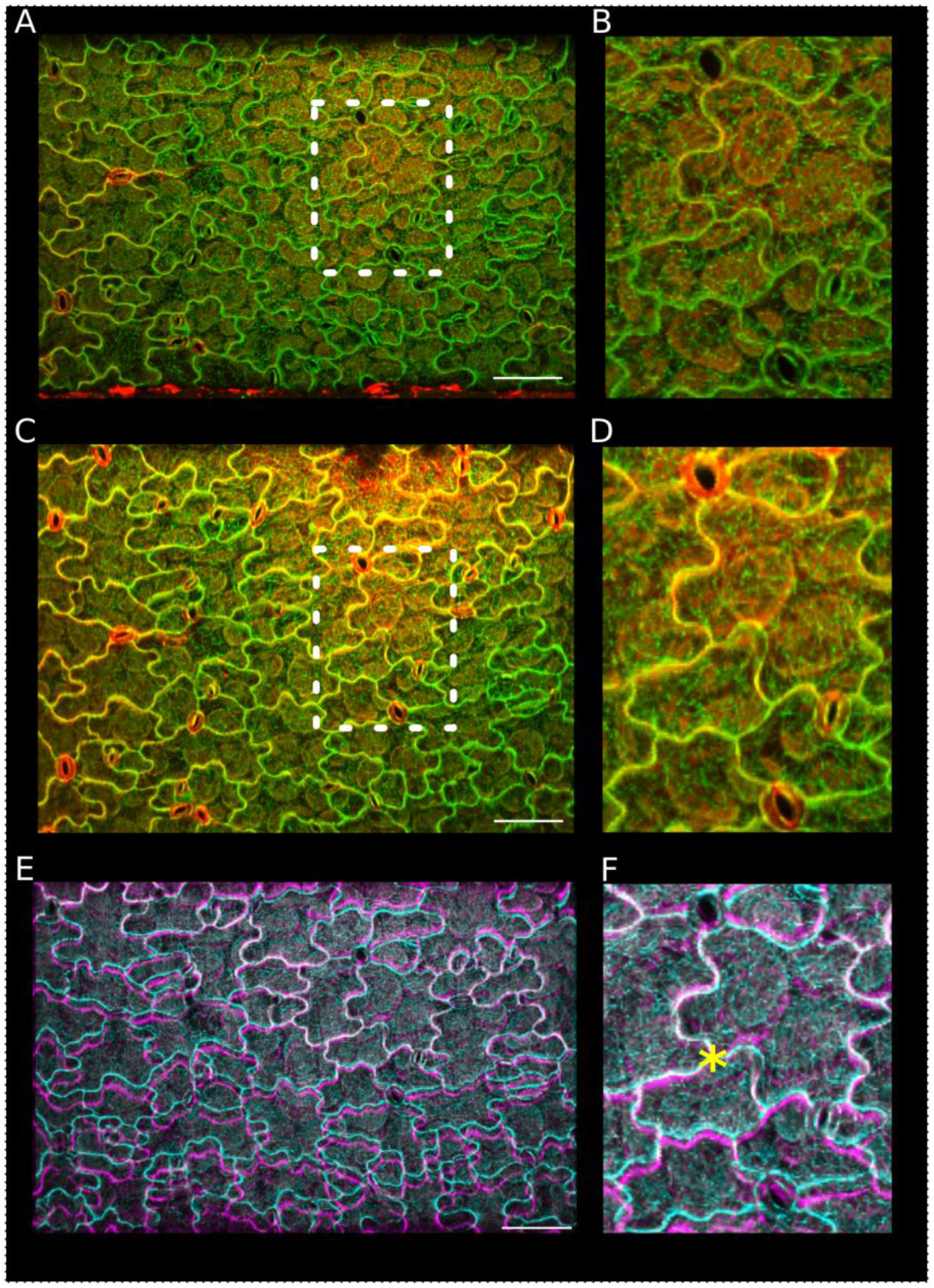
Using ACME to apply to stress to cotyledons. A cotyledon was mounted onto ACME. The sample expresses the MBD-GFP marker (green) was stained with PI (red) to demonstrate that the cells are healthy. **(A-B)** Image of the cotyledon obtained with the 20x objective before application of 5mN of stress, including a zoomed in area **(B)**. **(C-D)** After application of stress, including zoomed in panel **(D)**. The sample remains healthy, the cells are turgid, they continue to express GFP and PI does not enter the cells. **(E-F)** Visually the before and after images appear very similar, however, by overlaying the GFP (before-cyan, after – magenta) we can see the deformation. The images were aligned at an arbitrary point (*) in the center of the image using MorphoGraphX. Opacity was adjusted to enable both images to be seen. The cells in the after image are elongated in the y axis compared to the before image.

## Discussion

ACME is a versatile tool for quantifying mechanical properties in small tissues. Its reliance on confocal imaging and image analysis tools for computing mechanical properties make it more accessible to biologists who are typically more familiar with imaging techniques than mechanics. By modifying the base plate, ACME can be used in conjunction with a range of microscopes, enabling information to be obtained at different resolutions. We have demonstrated here the usefulness of ACME by measuring mechanical properties that could not previously be measured including: elasticity and creep in light grown Arabidopsis seedlings and revealed cellular resolution spatial gradients in mechanical properties that correlated with GA induced growth.

The layers of the hypocotyl are connected so the strain measured on the epidermis is equal to the strain across all of the layers. The strain observed reflects the properties of the whole tissue and will be most influenced by the load bearing layer of the tissue. The strain measured in the epidermis, therefore, provides a means of measuring the properties of the load bearing layer, even without knowing the identity of the layer. While the strain in length is equal across the cross section of the hypocotyl the change in diameter need not be and could provide information about the properties of the internal cell layers.

In addition to measuring mechanical properties there is a great interest in the community to look at mechanical feedback. A range of methods have been employed to investigate feedback regulation by mechanical stress on development. These include, making cuts or ablations (Hamant et al., 2008; Sampathkumar et al., 2014), osmotic stress (Nakayama et al., 2012), compressing the tissue (Nakayama et al., 2012; Sampathkumar et al., 2014), or large scale deformations (Bringmann and Bergmann, 2017). Taken together these methods suggest that plants do sense mechanical stress. However, all the methods also induce other types of stress to the tissue, for example wounding or drought. In many cases the exact stress being applied is unclear and has to be deduced from models. If the details of this response are to be worked out there is clearly a need for methods that enable more subtle application of quantifiable stress in a less invasive manner. ACME provides the opportunity to do this as we are able to continuously monitor the sample.

## Materials and Methods

### ACME Hardware

The extensometer is based on two SLC1720s nanopositioners (Figure 1A-C, label 4-5) (SmarAct GmbH). The positioners are controlled by a SmarAct MCS3D (SmarAct GmbH) controller (Figure 1C, label 15) accompanied by its software library, which in turn is controlled by custom-made software (available here: https://github.com/ACME-Robinson/InstallPackage) (Figure 1C, label 14). One of the positioners is driving the moving plate of the extensometer responsible for exerting force upon the sample (Figure 3, label 5), while the other positioner (Figure 1A-C, label 4) is responsible for the vertical motion of the main extensometer body to allow replacement of Petri dishes. The force measurement system consists of force sensor (Futek LSB200 10g load cell, Futek Inc.) (Figure 3A-C, label 3) its dedicated amplifier (Futek CSG110, Futek Inc.) (Figure 1C, label 11) and a computer-based signal acquisition system (NI6221PCI (Figure 1C, label 13) with SCB68 break-out box (Figure 1C, label 12), National Instruments). The force sensor is attached to the sensing plate (Figure 1A, C, label 2) of the extensometer. In contrast to the moving plate (Figure 1A-C, label 1), the sensing plate is not actuated, instead it is attached to a force sensor that measures force exerted on the plate by the sample. The load cell amplifier drives the load cell and picks up the force signal from the sensor. Amplified force signal from the amplifier’s output is delivered to the signal acquisition card for sampling. The load cell amplifier is wrapped in insulating foam to stabilize its temperature and therefore to minimize drift. The signal acquisition card mounted inside of a Linux PC is controlled by comedi library (linux control and measurement device interface). The comedi library provides software calibration solution utilizing internal calibration reference of the NI6221PCI card. The force signal is sampled at 10kHz frequency, software-calibrated and then averaged in a 200ms window (average of 2000 voltage samples). Then, the averaged voltage is offset-corrected (tare) with a value obtained during sensor tare at the beginning of the experiment. The offset-corrected voltage is converted into force by multiplying it by a gain factor [μN/V] obtained during calibration of the force measurement system. The obtained force F[μN] is therefore F = g(V – Vo), where g is the gain [μN/V], V is the measured voltage [V] and Vo is the offset voltage [V]. The MCS Control Library which controls the smarAct nan-positioners is supplied by the manufacturer and requires Linux for 32bit/x86 architecture. The extensometer plates (Figure 1A-C, label 1-2) as well as all its other components (Figure 1A-C label 8, and Figure 1C label 10) are custom-made out of readily available aluminium profiles and assembled together with standard screws and nuts. Detailed plans on assemble of ACME is included in the supplemental file (building_acme).

### ACME control software

The hardware is directly controlled by accompanying libraries which we interface through the custom-made ACMErobotX software (Figure 1C, label 14). We also developed the ACME software to perform higher level functions (both software are available here: https://github.com/ACME-Robinson/InstallPackage). To aid the user, we added a parameter file where the user can easily specify the key features of the experiment without any knowledge of coding. The parameter file is where the user can specify other features for example the speed of movement of the robot, the initial gap size, or the location to save the data. To further aid the user the robot position and measured force is saved continuously as a csv file and can be viewed live or analysed later.

### ACME calibration

The force sensor (load cell) is calibrated by detaching it from the extensometer and vertically mounting it on a stand. Weights are placed on top of the load cell in a sequence and their corresponding digitized voltages are acquired, as seen by the software. Least squares linear fitting provides the slope of the voltage-to-force relationship which we call sensor gain, expressed in μN/V. The gain value is later used by the software to compute force in μN from the acquired voltage values (ACMECalibrationGuide.pdf calibration_worksheet.odt). The aim of the calibration procedure is to determine the overall gain (scaling) of the force measurement system. The input is a series of weight readings of test weights measured with a precise laboratory scales. This information is put into the robot.ini file before the first use of the equipment. Drift was measured by holding the position of the plates and measuring the force. 51 runs were performed sequentially. It was found that drift was highest in the first run but after this it dropped to below 100μN per hour (Figure S1). This is 10% of the smallest force typically applied. Dimensional changes to the sample result in varying force being exerted on the extensometer plates (approximately 1.7μm for 1000μN).

### ACME protocols

Parameters have the following form: Force or position F/P instructs the system to move to achieve a target force or a target position. The position is relative to the previous position while the force is compared to the zero force set at the start of the experiment. The magnitude of force or the position is then given, position is in μm and force is in μN. The length of time to hold the new force or position is then given in seconds. At all times the force and position is being recorded. The plates continue to be adjusted to maintain the force if it goes outside the threshold specified. Usually a threshold of 100μN is used as this is comparable to the drift that we observed. The step size is also specified and can be altered depending on the size of the deformation or the property of the material. Usually a step size of 2μm is used. The initial gap size is also specified, this is how far apart the plates are at the start of the experiment.

Example Creep protocol:

The following example is of a creep test. The plates move to achieve zero force which is held for 1800 seconds, then the plant is stretched until a force of 1000 μN is exerted on the load cell, this is held for 1800 seconds. The force is then returned to 0.

Step1, F, 0, 1800
Step2, F, -1000, 1800
Step3, F, 0, 1800

#### Oscillation experiments

Samples were rapidly loaded and unloaded. Samples were held at the stated force for 1 second. 5-10 oscillations were made at each force then the sample was held at 0 force for 30 seconds then a new batch of oscillations was performed. Bright-field images were collected every 645ms. The images were opened in the ACMEtracker (software available here: https://github.com/ACME-Robinson/InstallPackage) and 2 regions of interest were selected. These regions were tracked in subsequent images using a normalized cross correlation coefficient method. The coordinates were written to a file and used to compute strain per batch of oscillations using R scripts (ExtractOsc.R). For 2DAS oscillation experiments at least 7 samples were tested per treatment, for 3DAS and uniconazole at least 5 samples were tested. The data from the replicates was combined using the stat_smooth function of ggplot2 in R.

To test competing hypothesis of a linear slope vs. a Hill function for the strain force relationship in the oscillation experiments we fitted both models using a least square estimate. It was not possible to apply as high a force to the 2DAS sample so the model was also fitted to a subset of the 3DAS data to match the range for the 2DAS samples. The force and strain data for the experiments was separated by treatment and age and fitted using the nonlinear least square method in R (nls). The linear models had the following form: ε=β.F + c, where ε is strain, F is force and, with starting parameters of β=0.1, c=0. The Hill function has the form:ε =α.F.(λ+F) -1 +0, with starting parameters α=0.1, λ=1000. The fitted values and residual sum of squares can be found in Figure 4- figure supplement 2 TableS1. As both models have 2 free parameters we can compare the RSS values to find the model that best fits the data.

#### Live creep tests

Live samples were mounted without glue and held at 0 force for 30 minutes then at 1mN for 3 hours then returned to 0 force for 30 minutes. Confocal z-stacks were collected regularly. A projection was made from the stack in MorphoGraphX and exported. Cells of interest were tracked using the Point tracker as in (Kuchen et al., 2012). The strain was computed manually form these measurements. Cells were selected that were as far apart as possible and vertically aligned. For the comparison of live, plasmolysed or frozen tissue, samples were frozen then thawed, alive in water or treated with 0.5M mannitol. For each treatment 3 samples were tested, with data points acquired every 1.5 minutes. For the creep tests on live samples 8 seedlings were used for GA and control conditions and 4 seedlings for the uniconazole condition.

#### Cellular resolution creep tests

Samples were held at 0 force for 15 minutes, then 10mN for 30 minutes then returned to 0 force for 20 minutes. Confocal stacks were collected and segmented as described for the growth measurements. Corresponding cells were then identified in subsequent images and given the same label. The change in cell size was computed to give cellular strain values. All cells that could be segmented in the images were used. Samples which detached or died were excluded. The gradient in cell properties in the 3DAS GA seedlings was seen in 4 samples, 1 sample did not show the gradient as it was curved.

#### ACME data analysis

The force and position are recorded to a csv file with the following headings: LoggerTime(ms), X_Position(um), Y_Position(um), IndentationForceSensor_Force(uN), IndentationForceSensor_OffsetVoltage(V), IndentationForceSensor_OffsetForce(uN), IndentationForceSensor_Voltage(V), ProtocolFlag(flag), SequenceStep. The first column records the time, the second and third records the position of the axis. The protocol flag can be used to separate the different stages of the experiment, notably the calibration step has a flag of 100 to distinguish it from the experiment itself. The other columns are only useful when calibrating the device for the first time, see supplemental information.

### Plant material, growth conditions and imaging

Seeds were surface sterilized and sown on MS medium [4.4gL-1 Murashige and Skoog salts (Sigma), 0.5 gL-1 4-morpholineethanesulphonic acid (Sigma), 0.8% agar, pH 5.7], with the possible addition of 10μΜ GA3, or 2μM uniconazole (Sigma 48880, 19701). Plants grown in the presence of uniconazole were germinated on control media and then transferred to the uniconazole after 2 days. Plants were imbibed in the dark at 4⁰C for 2 or 3 days then grown in controlled environment chambers (100μE continuous light conditions at 22⁰C) as in (Sauret-Güeto et al., 2012). *35S::PIN1-GFP* line from the Benkova lab and *35S::GFP-MBD* line (Hamant et al. 2008), was used to label cells. Cell walls were stained with 0.1% propidium iodide.

Images were acquired using a LEICA SP5 with a HCX APO L 20x/0.5 W objective and a Leica HyD hybrid detector. GFP images were obtained using a 488nm laser. The power was maintained as low as possible. For oscillation experiments images were collected in a single z-plane using the bright field detector, for rapid imaging. For growth curves, and creep tests Z-stacks were collected. To enable cell segmentation in MorphoGraphX a Z-step of 0.4-0.5μm was used, with a scan speed of 400Hz and no averaging.

### Growth analysis

Hypocotyl lengths were measured in ImageJ. Cell volume and length calculations were made using a version of MorphoGraphX (http://www.lithographx.org/) which contains the necessary processes. The image stacks were loaded into LithoGraphX and segmented by the following procedure. A gaussian blur of size (1, 1, 1), autoscaling of the stack, ITK autoseed segmentation with a threshold between 1000 and 1500 depending on the sample. Any over segmented cells were then merged, under segmented areas were deleted as were cells from the internal layers, those that were not complete, or those that were part of the stomata lineage as their size does not reflect that of the growth rate of the plant, but is under different regulation. Marching cubes of size 2 was performed without smoothing. A Bézier curve of order 5 was fitted to the sample, arc-length parameterized, with the zero set to a defined point along the hypocotyl. The position of a cell along the hypocotyl is then defined as the curve parameter of the projection of its center onto the Bézier curve. For each cell, we have computed its volume, length and mean radius. The volume could be estimated directly from the image. The length and mean radius was computed by first extracting the principal components of the cell shape. The main one correspond to the direction of the length and 2nd and 3rd to the width and height. The distances along a PC was measured by projecting the voxels defining a cell onto the direction and taking the distance between the 0.5 and the 99.5 percentiles. The mean radius is then the average of the lengths along the 2nd and 3rd axis. Any cells not well represented were deleted. Graphs were produced using ggplot in R; the stat smooth function was used to fit a confidence interval with 99% level. Local polynomial regression fitting was performed on the cell length data using the loess function in R. The output of this fitting was used to compute the difference in length of cells at the different time points.

### Sample attachment

Entire seedlings were attached without crushing the tissue (Sample_mounting.avi). The experiments were conducted in solution to prevent sample drying. The attachment of the sample to the movement arm was made using tape (0.94X0.50 inches, white, catalog no. TTSW-1000, DiversifiedBiotech). They are water proof and remain attached to the robot arm, even when immersed in solution. Samples remained healthy for many hours (longer experiments were not conducted) and can be tested with forces in the 1-2mN range. For higher force measurements (≥5mN) a thin layer of cyanoacrylate glue was added to the tape (glue.avi), under this condition the sample remains healthy for shorter periods of time. Sample health was assessed visually; samples were regarded as healthy if plasmolysis did not occur. Sample health could also be measured by monitoring the force, a healthy sample will exert a force if held at a fixed position and will expand if actively maintained at zero force. Cyanoacrylate glue is suitable for previously frozen tissue or for short experiments with high force.

#### AFM sample preparation and analysis

Plants used for AFM experiments were grown as described above except in long day conditions. Immediately preceding AFM analysis, each batch of seedlings was dissected, on moist paper towels, to remove the cotyledons. Dissected hypocotyls were then placed on etched glass slides and secured between glass bumpers with 0.8% low melt agarose in 0.55M manitol. Once secured, the samples were flooded with 0.55M mannitol to suppress turgor pressure. Samples were left to plasmolyze for a minimum of 20 minutes. All solutions were prepared using ultrapure water at pH 7.1.

Dissected and plasmolyzed hypocotyls were indented using a Nano Wizard 3 AFM (JPK Instruments, DE) mounted with a 0.8μm diameter rounded indenter (Windsor Scientific, UK) on a cantilever of 40 N/m stiffness. Cantilever stiffness was determined by thermal tuning prior to experiment initiation. Tip sensitivity was calibrated by first performing indentations on a clean glass slide. For each hypocotyl, three areas of 50x100μm were indented with 32x32 points: a top area near the cotyledons, a middle area, and an area just before the collet. Positions of each grid were recorded with a top view camera. Indentations were performed with 1000nN of force yielding an indentation depth range of 250-500nm and a total of 1024 indentations were performed per area per biological sample.

#### AFM data analysis

Force indentation curves were analyzed using JPKSPM Data Processing software (JPK Instruments, DE; v. spm 4.3.10) using the following steps: voltage reading were converted to for using calibrated sensitivity and cantilever stiffness values, baseline subtraction and tilt correction, vertical displacement offset adjustment, indentation calculation by subtraction of cantilever bending from piezo position during indentation, and elastic stiffness was calculated by fitting a tangent to the final 150 nm of the indentation. Extraction curves were not analyzed due to numerous adhesion difficulties during tip removal from the surface. Fitting the last 150nm of the indentation ensured that stiffness values reflected a linear elastic constant and avoided possible contact area evolution during the beginning of the indentation. Elastic stiffness maps were then imported into MatLab and values were selected from anticlinal cell walls only. Anticlinal walls were used for the following reasons: often surface walls buckled or showed complex force-deformation curves due to geometrical instability without turgor support, anticlinal walls maintained relatively constant depth during cell elongation allowing for comparability across cell sizes, and lastly anticlinal wall indentations were more normal to the indenter axis than curved surface walls. For each grid area, 30-50 points were chosen from anticlinal walls and used for subsequent analyses, representing data from 3-10 cells depending on cell length in the scan area.

#### Cell wall thickness measurements

Seedlings were fixed in 50 mM Na-caodylate buffer pH=7.4 with 2% (v/v) glutaraldehyde (EMS; http://www.ems-group.com) for 2h at room temperature. After 6 washes with 50 mM Na-caodylate buffer pH=7.4, the samples were postfixed overnight with 1% (w/v) OsO4 in Na-cacodylate buffer pH=7.4 at 4°C. After 6 washes in calcodylate buffer and 1 wash with water, the samples were dehydrated through an aceton series (10%, 20%, 30%, 50%, 70%, 90%, 100%, each 10 min), and seven subsequent changes of aceton. Embedding proceded by soaking in increasing concentrations of Spurr’s resin (Plano; https://www.plano-em.de) in aceton (25% 1.5h, 50% 1.5h, 75% overnight, 100% 6h), and polymerisation at 70°C for 18h under dry atmosphere (silicagel). Ultrathin sections (70-80 nm) were contrasted with 2% (w/v) uranyl acetate and subsequently with 80 mM lead citrate (Reynolds, 1963). Cell wall measurements were made in ImageJ. Multiple measurements were made per slice and per cell so calculate the average cell wall width. For each treatment sections were taken from two plants.

## Acknowledgement

We thank Prashant Saxena and Hagen Reinhardt for helpful discussions.

---

**Figure S1.**
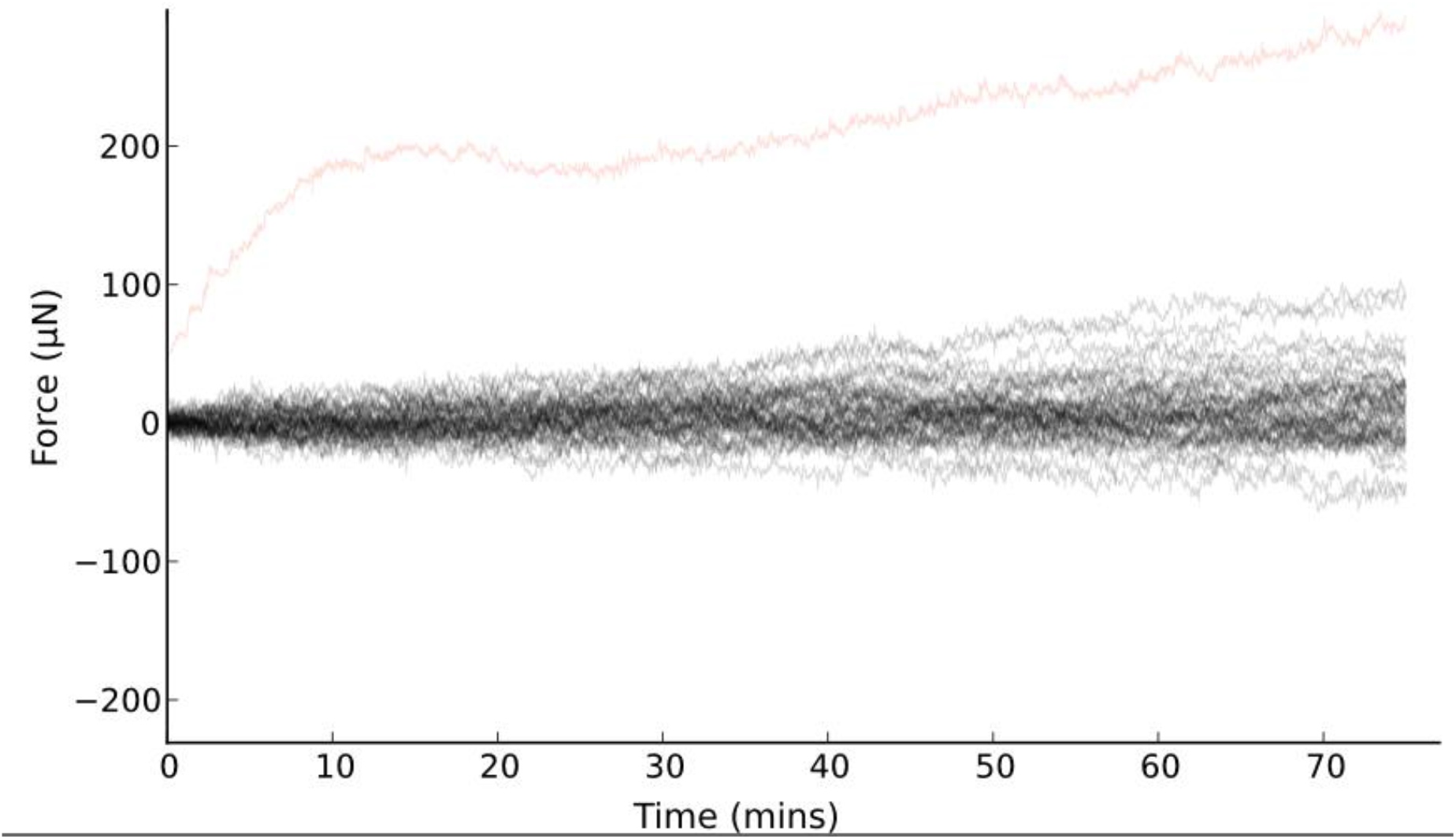
Drift assessment of ACME. Drift was measured by holding the position of the plates and measuring the force. 51 runs were performed sequentially. It was found that drift was highest in the first run (red) but after this it dropped to below 100μN per hour. This is 10% of the smallest force typically applied.

**Figure S2.**
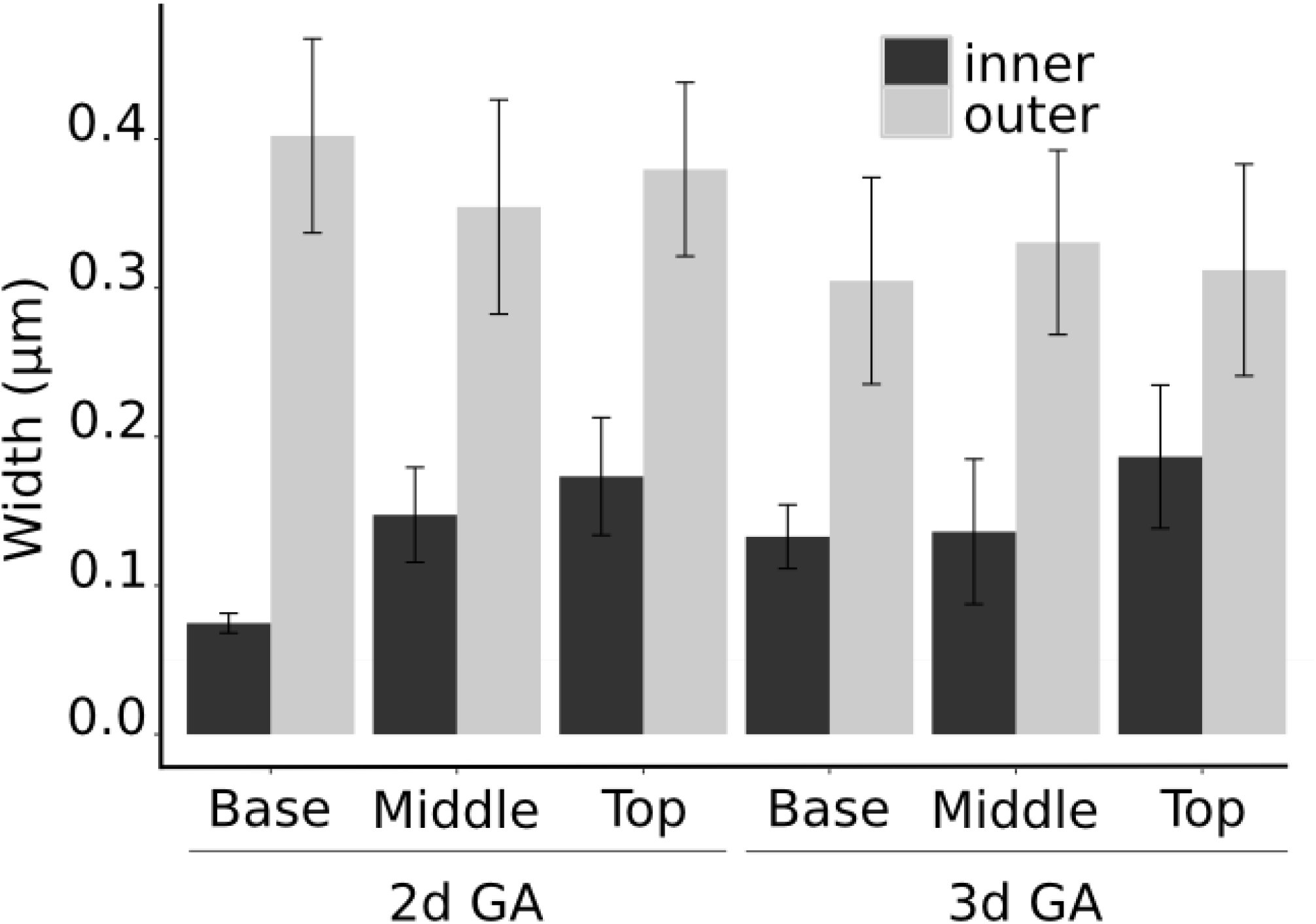
Cell wall width measured in ImageJ from TEM images at 3 locations along 2 and 3DAS GA treated seedlings. The inner and outer wall of the epidermal cells was measured. (Error bars show standard deviation, scale bars =100μm.)

## References

Adams, P.A., Montague, M.J., Tepfer, M., Rayle, D.L., Ikuma, H., and Kaufman, P.B. (1975). Effect of gibberellic Acid on the plasticity and elasticity of Avena stem segments. Plant Physiol. 56: 757–760.

Band, L.R. et al. (2012). Root gravitropism is regulated by a transient lateral auxin gradient controlled by a tipping-point mechanism. Proc. Natl. Acad. Sci. 109: 4668–4673.

Bastien, R., Bohr, T., Moulia, B., and Douady, S. (2013). Unifying model of shoot gravitropism reveals proprioception as a central feature of posture control in plants. Proc. Natl. Acad. Sci. 110: 755–760.

Braam, J. (2005). In touch - plant responses to mechanical stimuli. New Phytologist. 165:14698137.

Braybrook, S.A. and Peaucelle, A. (2013). Mechano-Chemical Aspects of Organ Formation in Arabidopsis thaliana: The Relationship between Auxin and Pectin. PLoS One 8: e57813.

Bringmann, M. and Bergmann, D.C. (2017). Tissue-wide Mechanical Forces Influence the Polarity of Stomatal Stem Cells in Arabidopsis. Curr. Biol. 27: 877–883.

Chehab, E.W., Eich, E., and Braam, J. (2009). Thigmomorphogenesis: A complex plant response to mechano-stimulation. J. Exp. Bot. 60: 43–56.

Coen E. and Rebocho A.B. (2016). Resolving Conflicts: Modeling Genetic Control of Plant Morphogenesis. Dev. Cell 38: 579–583.

Coen, E., Rolland-Lagan, A.-G., Matthews, M., Bangham, J.A., and Prusinkiewicz, P. (2004). The genetics of geometry. Proc. Natl. Acad. Sci. U. S. A. 101: 4728–4735.

Cosgrove, D.J. (2015). Plant cell wall extensibility: Connecting plant cell growth with cell wall structure, mechanics, and the action of wall-modifying enzymes. J. Exp. Bot. 67: 463–476.

Cosgrove, D.J. and Sovonick-Dunford, S. A (1989). Mechanism of gibberellin-dependent stem elongation in peas. Plant Physiol. 89: 184–91.

Ditengou, F.A., Teale, W.D., Kochersperger, P., Flittner, K.A., Kneuper, I., van der Graaff, E., Nziengui, H., Pinosa, F., Li, X., Nitschke, R., Laux, T., and Palme, K. (2008). Mechanical induction of lateral root initiation in Arabidopsis thaliana. Proc. Natl. Acad. Sci. U.S.A. 105: 18818–18823.

Dumais, J. and Steele, C. (2000). New Evidence for the Role of Mechanical Forces in the Shoot Apical Meristem. J. Plant Growth Regul. 19: 7–18.

Durachko, D.M. and Cosgrove, D.J. (2009). Measuring plant cell wall extension (creep) induced by acidic pH and by alpha-expansin. J. Vis. Exp. 25: 1263–1267.

Eldridge, T., Łangowski, Ł., Stacey, N., Jantzen, F., Moubayidin, L., Sicard, A., Southam, P., Kennaway, R., Lenhard, M., Coen, E.S., and Østergaard, L. (2016). Fruit shape diversity in the Brassicaceae is generated by varying patterns of anisotropy. Development 1: 3394–3406.

Fleming, A. J. (1997). Induction of Leaf Primordia by the Cell Wall Protein Expansin. Science 276: 1415–1418.

Fung, Y.-C. (1993). Biomechanics 2nd ed. (Springer New York: New York, NY).

Green, A.A., Kennaway, J.R., Hanna, A.I., Andrew Bangham, J., and Coen, E. (2010). Genetic control of organ shape and tissue polarity. PLoS Biol. 8: e1000537.

Green, P.B. (1999). Expression of pattern in plants: Combining molecular and calculus-based biophysical paradigms. Am. J. Bot. 86: 1059–1076.

Green, P., Steele, C., and Rennich, S. (1996). Phyllotactic patterns: a biophysical mechanism for their origin. Ann. Bot. 77: 515–528.

Green P.B. (1962). Mechanism for Plant Cellular Morphogenesis. Science 138: 1404–1405.

Hamant, O., Heisler, M.G., Jönsson, H., Krupinski, P., Uyttewaal, M., Bokov, P., Corson, F., Sahlin, P., Boudaoud, A., Meyerowitz, E.M., Couder, Y., and Traas, J. (2008). Developmental patterning by mechanical signals in Arabidopsis. Science 322: 1650–1655.

Hejnowicz, Z, and Sievers, A. (1995). Tissue stresses in organs of herbaceous plants II. Determination in three dimensions in the hypocotyl of sunflower. J. Exp. Bot. 46: 1045–1053.

Kierzkowski, D., Nakayama, N., Routier-Kierzkowska, A.-L., Weber, A., Bayer, E., Schorderet, M., Reinhardt, D., Kuhlemeier, C., and Smith, R.S. (2012). Elastic Domains Regulate Growth and Organogenesis in the Plant Shoot Apical Meristem. Science 335: 1096–1099.

Kuchen, E.E., Fox, S., Barbier de Reuille, P., Kennaway, R., Bensmihen, S., Avondo, J., Calder, G.M., Southam, P., Robinson, S., Bangham, A., and Coen, E. (2012). Generation of Leaf Shape Through Early Patterns of Growth and Tissue Polarity. Science 335: 1092–1096.

Kutschera, U. and Niklas, K.J. (2007). The epidermal-growth-control theory of stem elongation: An old and a new perspective. J. Plant Physiol. 164: 1395–1409.

Landau, L.D. and Lifshitz, E.M. (1986). Course of Theoretical Physics, vol. 7, Theory of Elasticity, Landau and Lifshitz. 7: 195.

Lipchinsky, A., Sharova, E.I., Medvedev, S.S., (2013). Elastic properties of the growth-controlling outer cell walls of maize coleoptile epidermis. Acta Physiol Plant 35:2183-2191.

Lockhart, J. (1965). An analysis of irreversible plant cell elongation. J. Theor. Biol. 8: 264–275.

Louveaux, M., Rochette, S., Beauzamy, L., Boudaoud, A. and Hamant, O. (2016), The impact of mechanical compression on cortical microtubules in Arabidopsis: a quantitative pipeline. Plant J, 88: 328–342.

McQueen-Mason, S., Durachko, D.M., and Cosgrove, D.J. (1992). Two endogenous proteins that induce cell wall extension in plants. Plant Cell 4: 1425–1433.

Miedes, E., Suslov, D., Vandenbussche, F., Kenobi, K., Ivakov, A., Van Der Straeten, D., Lorences, E.P., Mellerowicz, E.J., Verbelen, J.P., and Vissenberg, K. (2013). Xyloglucan endotransglucosylase/hydrolase (XTH) overexpression affects growth and cell wall mechanics in etiolated Arabidopsis hypocotyls. J. Exp. Bot. 64: 2481–2497.

Milani, P., Gholamirad, M., Traas, J., Arnéodo, A., Boudaoud, A., Argoul, F., and Hamant, O. (2011). In vivo analysis of local wall stiffness at the shoot apical meristem in Arabidopsis using atomic force microscopy. Plant J. 67: 1116–1123.

Milani, P., Mirabet, V., Cellier, C., Rozier, F., Hamant, O., Das, P., and Boudaoud, A. (2014). Matching Patterns of Gene Expression to Mechanical Stiffness at Cell Resolution through Quantitative Tandem Epifluorescence and Nanoindentation. Plant Physiology.165:1399-1408.

Moulia, B. et al. (2011). Integrative mechanobiology of growth and architectural development in changing mechanical environments. In Mechanical Integration of Plant Cells and Plants, pp. 269–302.

Nakayama, N., Smith, R.S., Mandel, T., Robinson, S., Kimura, S., Boudaoud, A., and Kuhlemeier, C. (2012). Mechanical regulation of auxin-mediated growth. Curr. Biol. 22: 1468–1476.

Paredez, A.R., Somerville, C.R., and Ehrhardt, D.W. (2006). Visualization of cellulose synthase demonstrates functional association with microtubules. Science 312: 1491–1495.

Park, Y.B. and Cosgrove, D.J. (2012). Changes in Cell Wall Biomechanical Properties in the Xyloglucan-Deficient xxt1/xxt2 Mutant of Arabidopsis. Plant Physiol. 158: 465–475.

Peaucelle, A., Braybrook, S. A., Le Guillou, L., Bron, E., Kuhlemeier, C., and Höfte, H. (2011). Pectin–induced changes in cell wall mechanics underlie organ initiation in Arabidopsis. Curr. Biol. 21: 1720–1726.

Peaucelle, A., Louvet, R., Johansen, J.N., Höfte, H., Laufs, P., Pelloux, J., and Mouille, G. (2008). Arabidopsis Phyllotaxis Is Controlled by the Methyl-Esterification Status of Cell-Wall Pectins. Curr. Biol. 18: 1943–1948.

Peters, W.S. and Tomos, A D. (1996). The history of tissue tension. Ann. Bot. 77: 657–65.

Pien, S., Wyrzykowska, J., and Fleming, A J. (2001). Novel marker genes for early leaf development indicate spatial regulation of carbohydrate metabolism within the apical meristem. Plant J. 25: 663–674.

Probine, M.C. and Preston, R.D. (1961). Cell growth and the structure and mechanical properties of the wall in internodal cells of Nitella opaca: I. Wall Structure and Growth. J. Exp. Bot. 12: 261–282.

Richter, G.L., Monshausen, G.B., Krol, A., and Gilroy, S. (2009). Mechanical stimuli modulate lateral root organogenesis. Plant Physiol. 151: 1855–1866.

Routier-Kierzkowska, A.-L., Weber, A., Kochova, P., Felekis, D., Nelson, B.J., Kuhlemeier, C., and Smith, R.S. (2012). Cellular Force Microscopy for in Vivo Measurements of Plant Tissue Mechanics. Plant Physiol. 158: 1514–1522.

Sampathkumar, A., Krupinski, P., Wightman, R., Milani, P., Berquand, A., Boudaoud, A., Hamant, O., Jönsson, H., and Meyerowitz, E.M. (2014). Subcellular and supracellular mechanical stress prescribes cytoskeleton behavior in Arabidopsis cotyledon pavement cells. Elife 3: e01967.

Sauret-Güeto, S., Calder, G., and Harberd, N.P. (2012). Transient gibberellin application promotes Arabidopsis thaliana hypocotyl cell elongation without maintaining transverse orientation of microtubules on the outer tangential wall of epidermal cells. Plant J. 69: 628–639.

Stuart, D. a and Jones, R.L. (1977). Roles of Extensibility and Turgor in Gibberellin-and Dark-stimulated Growth. Plant Physiol. 59: 61–68.

Weber, A., Braybrook, S., Huflejt, M., Mosca, G., Routier-Kierzkowska, A.L., and Smith, R.S. (2015). Measuring the mechanical properties of plant cells by combining micro-indentation with osmotic treatments. J. Exp. Bot. 66: 3229–3241.

